# Socioeconomic status of indigenous peoples with active tuberculosis in Brazil: a principal components analysis

**DOI:** 10.1101/290668

**Authors:** Laís P. Freitas, Reinaldo Souza-Santos, Ida V. Kolte, Jocieli Malacarne, Paulo C. Basta

**Affiliations:** National School of Public Health Sergio Arouca. Oswaldo Cruz Foundation (FIOCRUZ). Rua Leopoldo Bulhões 1480, Manguinhos, Rio de Janeiro/RJ, Brazil, 21041-210.

**Keywords:** INDIGENOUS PEOPLE, POVERTY, TUBERCULOSIS, SOCIOECONOMIC FACTORS, SOUTH AMERICAN INDIANS

## Abstract

Indigenous people usually live in precarious conditions and suffer a disproportionally burden of tuberculosis in Brazil. To characterize the socioeconomic status of indigenous peoples with active tuberculosis in Brazil, this cross-sectional study included all Amerindians that started tuberculosis treatment between March 2011 and December 2012 in four municipalities of Mato Grosso do Sul state (Central-Western region). We tested the approach using principal components analysis (PCA) to create three socioeconomic indexes (SEI) using groups of variables: household characteristics, ownership of durable goods, and both. Cases were then classified into tertiles, with the 1^st^ tertile representing the most disadvantaged. A total of 166 indigenous cases of tuberculosis were included. 31.9% did not have durable goods. 25.9% had family bathroom, 9.0% piped water inside the house and 53.0% electricity, with higher proportions in Miranda and Aquidauana. Houses were predominantly made using natural materials in Amambai and Caarapó. Miranda and Aquidauana had more cases in the 3^rd^ tertile (92.3%) and Amambai, in the 1^st^ tertile (37.7%). The indexes showed similar results and consistency for socioeconomic characterization. The percentage of people in the 3^rd^ tertile increased with years of schooling. The majority in the 3^rd^ tertile received Bolsa Família, a social welfare programme. This study confirmed the applicability of the PCA using information on household characteristics and ownership of durable goods for socioeconomic characterization of indigenous groups and provided important evidence of the unfavorable living conditions of Amerindians with tuberculosis in Mato Grosso do Sul.

## INTRODUCTION

Tuberculosis is a major public health problem in Brazil, with approximately 70000 new cases and 4400 tuberculosis-related deaths per year.^1^ The disease mostly affects underprivileged groups such as indigenous people. Data from the Ministry of Health show that, in 2010, the incidence of tuberculosis among the indigenous population was two and a half times the average incidence of the country (94.9 and 37.6 per 100000 inhabitants, respectively).^2,3^ A recent study showed that the indigenous populations had the highest tuberculosis incidences in all Brazilian regions, except the South, with a greater difference to the incidence of other ethnic groups in the Central-West region.^4^ In this region, in the state of Mato Grosso do Sul during the period 2001 to 2009, 15.6% of notified tuberculosis cases were indigenous, although they represented only three percent of the population. The indigenous population had the highest incidence of all ethnic categories for all years studied, with a mean of 209.0 cases per 100000 inhabitants, more than six times the overall mean of the state (34.5 per 100000).^5^

Some factors commonly found among the indigenous population contribute to the high burden of tuberculosis. Indigenous peoples are among the most impoverished groups in the world as a consequence of historic injustice, colonization, dispossession of their lands, oppression and discrimination.^6^ Previous studies in the general population have established unfavorable socioeconomic conditions as a risk factor for tuberculosis.^7–10^ Despite accounting for nearly five per cent of the world’s population, indigenous peoples constitute 15% of the world’s poor and one third of the world’s 900 million extremely poor rural people.^6^ Data on the socioeconomic conditions of indigenous peoples in Brazil are scarce. The First National Survey of Indigenous People’s Health and Nutrition in Brazil (henceforth, “National Survey”), conducted between 2008 and 2009, represents a key effort to start filling this gap, obtaining information on the socioeconomic status and sanitary conditions.^11^

The socioeconomic characterization is a challenge, especially in vulnerable groups such as the indigenous peoples. Using income data requires exhaustive data collection. Also, income as an indicator often fails to capture whether people have income in kind and trade goods, and is difficult to measure for people with temporary jobs.^12,13^ An alternative indicator could be information on consumption or expenditure, but accurate and complete data are hard to obtain. Therefore, data on ownership of durable goods and household characteristics have been used for socioeconomic characterization as an alternative method with simpler data collection.^14^ This approach reflect long-term household wealth and living standards, which is important in the context of tuberculosis, considering its relationship with poverty and established association with possession of few goods.^7,15^

Given that indigenous peoples suffer a disproportionately high burden of tuberculosis and the scarcity of data on the living conditions of this population in Brazil, the aim of this study was to characterize the socioeconomic status of indigenous peoples with active tuberculosis in Brazil, applying a principal components analysis to generate and compare different socioeconomic indexes based on the ownership of durable goods and the household characteristics. To our knowledge, this is the first time that the applicability of such approach is tested for indigenous groups.

## MATERIALS AND METHODS

### Study area, population and design

This was a cross-sectional study including indigenous cases of active tuberculosis recently diagnosed by the healthcare system and who initiated treatment between March 2011 and December 2012 in one of the selected Polo-Bases of Mato Grosso do Sul (Aquidauana, Miranda, Caarapó and Amambai). The Polo-Base is an administrative unit responsible for primary healthcare under the Brazilian Indigenous Health System, a subsystem of the Unified Health System (SUS). A Polo-Base is usually located in key municipalities and has its own multidisciplinary indigenous healthcare team.^16,17^ Aquidauana and Miranda are located in the northern part of the state and were analyzed together due to the small numbers of indigenous cases of tuberculosis. Caarapó and Amambai are in the southern part of the state, close to the border with Paraguay. The largest ethnic groups living in these areas are the Guarani-Kaiowá and the Terena. Together, these Polo-Bases provide assistance for 35 villages and 33267 indigenous people (6984 in Aquidauana, 7217 in Miranda, 6150 in Caarapó and 12916 in Amambai), nearly half of the indigenous population of the state (71658).^18^

### Data collection

Participants (or participants’ parents, for minors) were interviewed by trained members of the indigenous healthcare team using a standard questionnaire. Information collected included age, sex, village, schooling, source of income, household characteristics and durable goods.

### Statistical analysis

All statistical analyses were performed using the software SPSS Statistics, version 20.0 (IBM, Armonk, NY, USA). Zero values were omitted from tables for better visualization of the data. The principal components analysis (PCA) was used to calculate socioeconomic indexes (SEIs). The PCA is a technique for dimensionality reduction that produces a smaller number of derived uncorrelated variables that can be used in place of a larger number of original correlated variables. It is increasingly used for socioeconomic characterization because this technique assigns weights to each variable included that represents the contribution of the given variable in the overall socioeconomic condition.^14^

To reach the best possible index with the collected data, three different PCAs were generated, based on the correlation matrix: 1) Goods PCA, including durable goods variables; 2) Household PCA, including household characteristics; and 3) Combined PCA, including both durable goods and household characteristics. All variables were first analyzed using descriptive statistics. Those with zero variance were not included in the PCA. Only durable goods available in five households or more were considered. Durable goods variables were organized as binary (0=No, 1=Yes). Household characteristics included continuous variables (number of dormitories and number of people sleeping in the same room), binary variables (family bathroom, i.e. of exclusive use of the family, and electricity) and categorical variables (type of floor, wall and roof). Recommendations from Kolenikov & Angeles were followed and categorical variables were organized as ordinals, with the lowest value corresponding to the most unfavorable socioeconomic condition and the highest value, to the most favorable condition.^19^ The results of the PCAs were compared with the Kaiser-Meyer-Olkin test to check if the model in use was properly fitted to the data (a minimum value of 0.600 is recommended).^20^

Given that three different PCAs were generated, each household came up with three different SEIs (Goods, Household, and Combined). Only the first component of each PCA was used, since it is the one described as related to the socioeconomic condition.^14^ Based on the first component, a SEI was calculated for each household through the sum of the contribution of each item (i.e. the “weight” obtained via PCA) times the value of the original variable. Next, households were classified in tertiles based on the SEI. The SEIs were compared to check for the most suitable index. Histograms of the SEIs were used to check for indication of clumping or truncation. Clumping occurs when households are grouped together in a small number of distinct clusters, and truncation, when the index is not able to differentiate between close socioeconomic groups easily (e.g. between the poor and the very poor). The means of each tertile for all three SEIs were calculated. Finally, the three SEIs classified in tertiles were cross-tabulated with variables that potentially have influence in the socioeconomic status (included or not in one or more PCAs) to check for consistency of the generated indexes. Bolsa Família, a social welfare programme of the Brazilian government that provides financial aid to poor families, was one of the explored variables.

### Ethical approval

The study was approved by the Brazilian National Committee for Ethics in Research (CONEP) and by the Ethics Committee of the National School of Public Health Sergio Arouca (ENSP), Fiocruz. Writen informed consent was obtained from all participants (or participants’ parents, for minors) prior to the interview.

## RESULTS

From March 2011 to December 2012, 168 indigenous people started anti-tuberculosis treatment in the study area. There was no refusal to participate, meaning all indigenous persons diagnosed with tuberculosis in the study area in the aforementioned period were included. The indigenous healthcare team later informed us that two of the participants were misdiagnosed with tuberculosis. These two patients were excluded from the analyses, resulting in a final study population of 166 indigenous cases of tuberculosis.

The majority (114, 68.7%) was treated in Amambai. Men predominated in all Polo-Bases, as well as adults aged 20-44 years (Table 1). The mean age was 38.0 years (range 1–87).

**Table 1.** Indigenous patients in treatment for tuberculosis by sex and age group, according to Polo-Base, Mato Grosso do Sul, Brazil, March 2011 to December 2012.

| CHARACTERISTIC |  | POLO-BASE |  |  |  |  |  | TOTAL |  |
| --- | --- | --- | --- | --- | --- | --- | --- | --- | --- |
|  |  | Miranda/<br>Aquidauana<br>(N=13) |  | Caarapó<br>(N=39) |  | Amambai<br>(N=114) |  | (N=166) |  |
|  |  | n | % | n | % | n | % | n | % |
| Sex | Male | 10 | 76.9 | 25 | 64.1 | 67 | 58.8 | 102 | 61.4 |
|  | Female | 3 | 23.1 | 14 | 35.9 | 47 | 41.2 | 64 | 38.6 |
| Age group<br>(years) | 0 to 4 | 0 | 0.0 | 4 | 10.3 | 2 | 1.8 | 6 | 3.6 |
|  | 5 to 14 | 0 | 0.0 | 4 | 10.3 | 5 | 4.4 | 9 | 5.4 |
|  | 15 to 19 | 1 | 7.7 | 1 | 2.6 | 5 | 4.4 | 7 | 4.2 |
|  | 20 to 44 | 10 | 76.9 | 16 | 41.0 | 67 | 58.8 | 93 | 56.0 |
|  | 45 to 64 | 2 | 15.4 | 8 | 20.5 | 15 | 13.2 | 25 | 15.1 |
|  | 65 or<br>more | 0 | 0.0 | 6 | 15.4 | 20 | 17.5 | 26 | 15.7 |

Overall, indigenous with tuberculosis from Miranda and Aquidauana proportionally had more durable goods. Fifty-three (31.9%) of the 166 indigenous declared having no durable goods in the household. Nobody declared owning an outboard motor. The use of materials from nature (e.g. straw, palm leaves) for building was more common in Caarapó and Amambai, while in Miranda and Aquidauana industrialized materials (e.g. bricks, cement) were mainly used (Table 2).

**Table 2.**
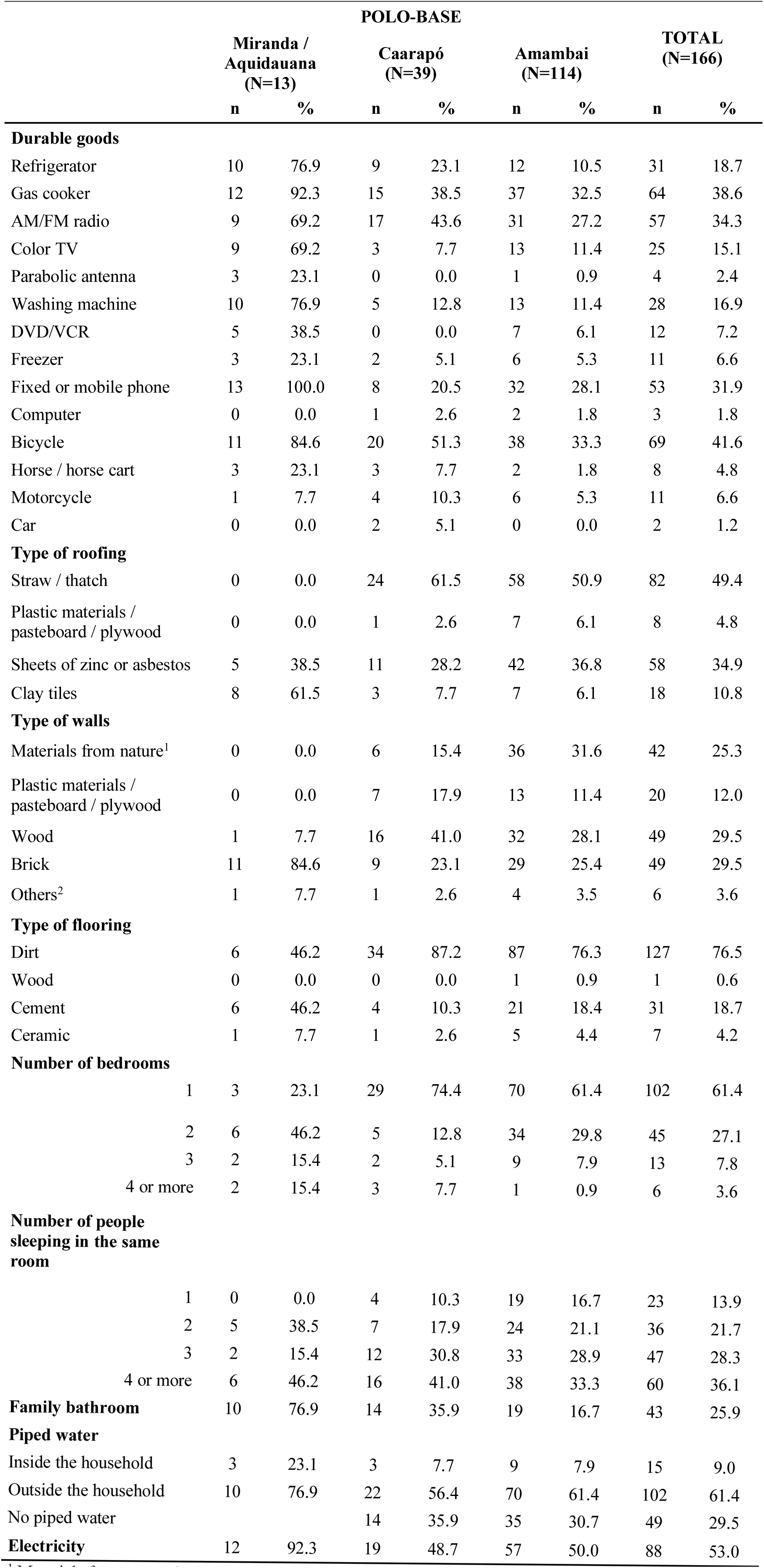
Characteristics of households of indigenous patients in treatment for tuberculosis according to Polo-Base, Mato Grosso do Sul, Brazil, March 2011 to December 2012.

Most households had only one bedroom (102/166 or 61.4%). In Miranda and Aquidauana more bedrooms per household were more common. The largest proportion (60/166 or 36.1%) reported sleeping with at least four other people. Only 25.9% (43/166) reported having a family bathroom. The proportions were consistently higher in Miranda and Aquidauana and lower in Amambai. Only 9.0% (15/166) of tuberculosis patients had piped water inside the household and 29.5% (43/166, all from Caarapó and Amambai) did not have a source of piped water. Almost half of households had electricity (88/166 or 53.0%), with higher proportions in Miranda and Aquidauana (Table 2).

Table 3 presents the contribution of each variable in the first component of each PCA. For the Goods PCA, socioeconomic status was more influenced by the ownership of a washing machine, a refrigerator, and a gas cooker. For the Household PCA, the types of roofing, walls, and flooring, as well as family bathroom, were the most influential variables. The negative influence of number of people sleeping in the same room indicates that the more people sleeping together, the lower the socioeconomic status. For the Combined PCA, types of coverage, walls, and flooring, gas cooker and washing machine remained among the most influential variables, while refrigerator and family bathroom diminished and electricity became more important. In general, the values of the contribution of each variable were close to those found separately for the Goods PCA and the Household PCA. The number of people sleeping in the same room maintained the negative association with socioeconomic status (Table 3).

**Table 3.** Component matrix of each Principal Components Analysis (PCA) performed.

| VARIABLES | Goods<br>PCA | Household<br>PCA | Combined<br>PCA |
| --- | --- | --- | --- |
| **Durable goods** |  |  |  |
| Refrigerator | 0.730 | NA | 0.689 |
| Gas cooker | 0.718 | NA | 0.717 |
| AM/FM radio | 0.605 | NA | 0.535 |
| Color TV | 0.660 | NA | 0.552 |
| Washing machine | 0.748 | NA | 0.704 |
| DVD/VCR | 0.616 | NA | 0.482 |
| Freezer | 0.444 | NA | 0.402 |
| Fixed or mobile phone | 0.688 | NA | 0.658 |
| Bicycle | 0.519 | NA | 0.434 |
| Horse / horse cart | 0.427 | NA | 0.312 |
| Motorcycle | 0.393 | NA | 0.392 |
| **Household characteristics** |  |  |  |
| Type of roofing | NA | 0.865 | 0.768 |
| Type of walls | NA | 0.816 | 0.743 |
| Type of flooring | NA | 0.789 | 0.705 |
| Number of dormitories | NA | 0.599 | 0.514 |
| Number of people sleeping in the same room | NA | -0.269 | -0.163 |
| Family bathroom | NA | 0.742 | 0.672 |
| Piped water | NA | 0.715 | 0.625 |
| Electricity | NA | 0.687 | 0.706 |

When comparing characteristics of the PCAs, the Household PCA presented a higher value in the Kaiser-Meyer-Olkin test (0.838 compared with 0.751 and 0.815 for the Goods PCA and the Combined PCA, respectively). The Household PCA also presented a higher percentage of variance explained by the variables (43.8% compared with 40.0% and 32.6% for the Goods PCA and the Combined PCA, respectively).

The distribution of households by each SEI is available in Figure 1. An example of truncation is seen in Figure 1A. The Goods SEI distribution presented an asymmetric distribution to the right and indicates that this index was not efficient in distinguishing households of lower socioeconomic conditions. This is a consequence of the high number of indigenous people that did not report ownership of any durable goods (Table 2). The Household SEI (Figure 1B) was closer to the normal distribution and showed little evidence of clumping or truncation, suggesting this index was better for distinguishing the socioeconomic condition. The Combined SEI (Figure 1C) appears to be between the Goods and the Household indexes, with a more even distribution and less evidence of clumping and truncation.

**Figure 1.**
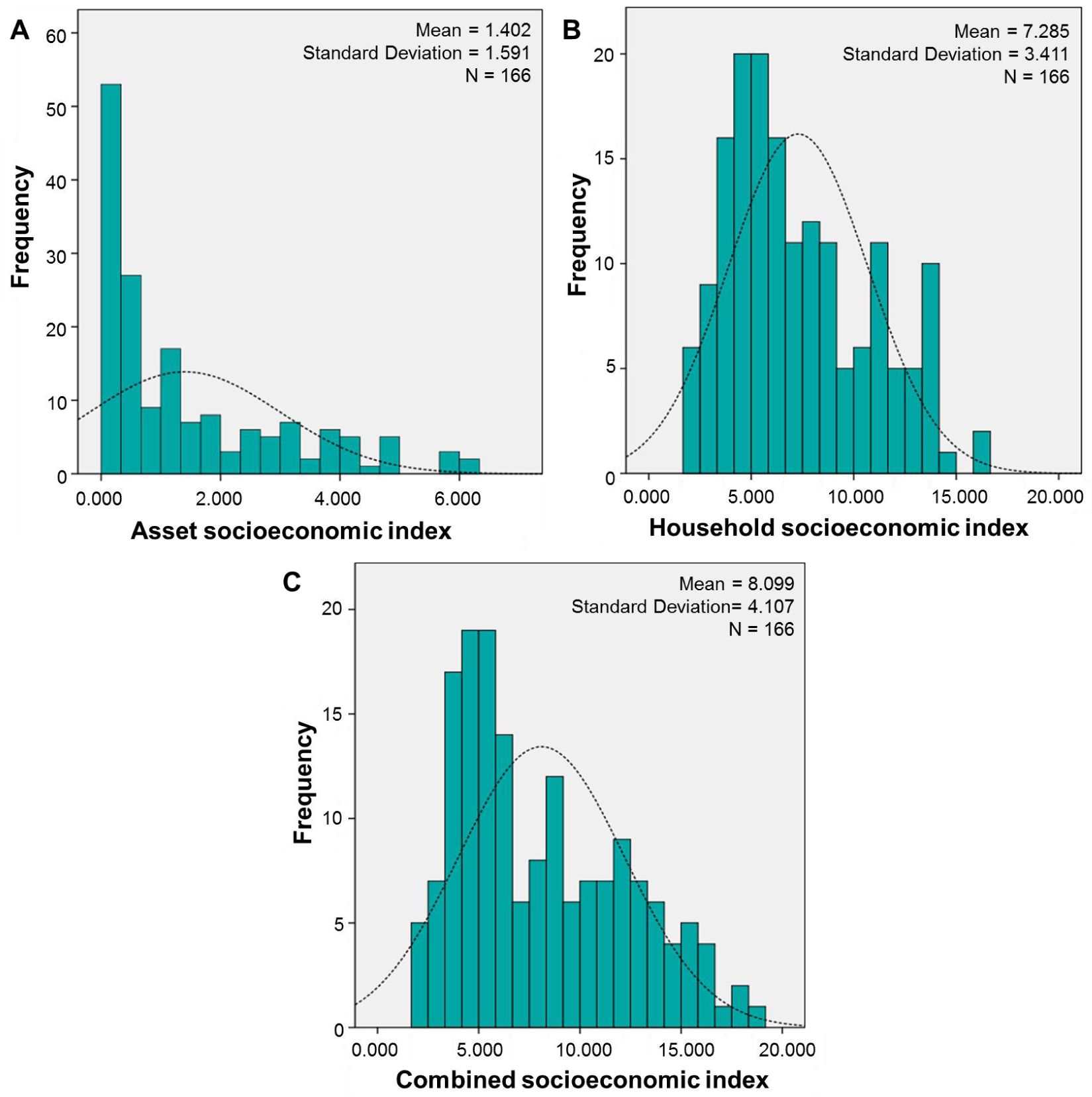
Distribution of households of indigenous people in treatment for tuberculosis by socioeconomic index (A: based on ownership of durable goods; B: based on characteristics of the household structure; C: based on both ownership of goods and characteristics of the household structure), Mato Grosso do Sul, Brazil, March 2011 to December 2012.

Comparing between the Polo-Bases, all indexes showed Miranda and Aquidauana with highest proportions of households in the 3rd tertile, and Amambai, in the 1st tertile (Table 4). Comparison of means inside the same tertile also revealed better socioeconomic conditions among indigenous people from Miranda and Aquidauana. These Polo-Bases had the highest means for every tertile, meaning a person from Miranda or Aquidauana had, on average, a better socioeconomic status than a person from Caarapó or Amambai classified in the same tertile. For Amambai, the Goods SEI mean was zero for the 1^st^ tertile, showing that all persons classified in this tertile in this Polo-Base did not have any durable goods in their households (Table 5).

**Table 4.** Indigenous patients in treatment for tuberculosis by tertile of socioeconomic index (SEI), according to Polo-Base, Mato Grosso do Sul, Brazil, March 2011 to December 2012.

|  | POLO-BASE |  |  |  |  |  |
| --- | --- | --- | --- | --- | --- | --- |
|  | Miranda/<br>Aquidauana<br>(N=13) |  | Caarapó<br>(N=39) |  | Amambai<br>(N=114) |  |
|  | n | % | n | % | n | % |
| **Goods SEI<sup>1</sup>** |  |  |  |  |  |  |
| 1<sup>st</sup> tertile | 0 | 0.0 | 11 | 28.2 | 43 | 37.7 |
| 2<sup>nd</sup> tertile | 1 | 7.7 | 13 | 33.3 | 43 | 37.7 |
| 3<sup>rd</sup> tertile | 12 | 92.3 | 15 | 38.5 | 28 | 24.6 |
| **Household SEI<sup>2</sup>** |  |  |  |  |  |  |
| 1<sup>st</sup> tertile | 0 | 0.0 | 13 | 33.3 | 43 | 37.7 |
| 2<sup>nd</sup> tertile | 3 | 23.1 | 16 | 41.0 | 36 | 31.6 |
| 3<sup>rd</sup> tertile | 10 | 76.9 | 10 | 25.6 | 35 | 30.7 |
| **Combined SEI<sup>3</sup>** |  |  |  |  |  |  |
| 1<sup>st</sup> tertile | 0 | 0.0 | 12 | 30.8 | 43 | 37.7 |
| 2<sup>nd</sup> tertile | 1 | 7.7 | 17 | 43.6 | 38 | 33.3 |
| 3<sup>rd</sup> tertile | 12 | 92.3 | 10 | 25.6 | 33 | 28.9 |

<sup>1</sup> Based on ownership of durable goods.
<sup>2</sup> Based on characteristics of the household structure.
<sup>3</sup> Based on both ownership of durable goods and characteristics of the household structure.

**Table 5.** Mean socioeconomic index (SEI) of households of indigenous people in tuberculosis treatment by tertile and Polo-Base, Mato Grosso do Sul, Brazil, March 2011 to

| POLO-BASE | Goods SEI<sup>1</sup> |  |  | Household SEI<sup>2</sup> |  |  | Combined SEI<sup>3</sup> |  |  |
| --- | --- | --- | --- | --- | --- | --- | --- | --- | --- |
|  | 1<sup>st</sup> tertile | 2<sup>nd</sup> tertile | 3<sup>rd</sup> tertile | 1<sup>st</sup> tertile | 2<sup>nd</sup> tertile | 3<sup>rd</sup> tertile | 1<sup>st</sup> tertile | 2<sup>nd</sup> tertile | 3<sup>rd</sup> tertile |
| Miranda/Aquidauana | 0.000 | 1.406 | 4.510 | 0.000 | 7.322 | 12.614 | 0.000 | 8.694 | 14.885 |
| Caarapó | 0.036 | 0.801 | 2.811 | 3.908 | 6.582 | 11.167 | 4.223 | 7.261 | 12.650 |
| Amambai | 0.000 | 0.850 | 3.135 | 3.866 | 6.567 | 11.168 | 3.949 | 7.079 | 12.658 |
| TOTAL | 0.007 | 0.848 | 3.346 | 3.875 | 6.612 | 11.431 | 4.009 | 7.163 | 13.142 |

<sup>1</sup> Based on ownership of durable goods.
<sup>2</sup> Based on characteristics of the household structure.
<sup>3</sup> Based on both ownership of durable goods and characteristics of the household structure.

Almost half of the population had no education (71/166 or 42.8%), 37.3% (62/166) began and/or completed the first four years of complementary school, 14.5% (24/166) began and/or completed the final four years of complementary school, four (2.4%) began high school, three (1.8%) finished high school and only two (1.2%) began college. For the three SEIs, people with no education were mostly classified in the 1^st^ and 2^nd^ tertiles (Goods SEI: 40.8% and 45.1%; Household SEI: 42.3% and 35.2%; Combined SEI: 38.0% and 43.7%, respectively). As years of schooling increased, the percentage of people in the 3^rd^ tertile increased as well, reaching 100.0% of those who finished high school and/or began college. Only eight people reported regular employment as a source of income. Among them, the majority was classified in the 3^rd^ tertile (Goods SEI: 62.5%; Household SEI and Combined SEI: 75.0%, each). The Goods SEI was the only index that classified at least one person with regular employment in the 1^st^ tertile. Bolsa Família was a source of income for 62/166 (37.3%) households. The majority of households in the 3rd tertile received Bolsa Família (Goods SEI: 33/55 or 60.0%; Household SEI: 28/55 or 50.9%; Combined SEI: 31/55 or 56.4%), while the majority classified in the 1^st^ tertile did not receive this financial aid (Goods SEI: 40/54 or 74.1%; Household SEI: 45/56 or 80.4%; Combined SEI: 45/55 or 81.8%).

A family bathroom was not available in nearly all households classified in the 1^st^ tertile, and available in the majority of those in the 3^rd^ tertile. All households in the 1^st^ tertile of the Household SEI and of the Combined SEI had dirt floors. All households with ceramic floors were classified in the 3^rd^ tertile by the three SEIs. Households with no source of piped water were more common in the 1^st^ tertile and least common in the 3^rd^ tertile. Those with piped water inside the house were mainly classified in the 3^rd^ tertile. The vast majority of households in the 1^st^ tertile did not have electricity, while in the 3^rd^ tertile, the majority did. Washing machine was mainly present in the 3^rd^ tertile of all three SEIs (Table 6).

**Table 6.** Characteristics of households of indigenous people in treatment for tuberculosis by tertile of socioeconomic index (SEI), Mato Grosso do Sul, Brazil, March 2011 to December 2012.

|  | Goods SEI<sup>1</sup> |  |  |  |  |  | Household SEI<sup>2</sup> |  |  |  |  |  | Combined SEI<sup>3</sup> |  |  |  |  |  |
| --- | --- | --- | --- | --- | --- | --- | --- | --- | --- | --- | --- | --- | --- | --- | --- | --- | --- | --- |
|  | 1<sup>st</sup> tertile |  | 2<sup>nd</sup> tertile |  | 3<sup>rd</sup> tertile |  | 1<sup>st</sup> tertile |  | 2<sup>nd</sup> tertile |  | 3<sup>rd</sup> tertile |  | 1<sup>st</sup> tertile |  | 2<sup>nd</sup> tertile |  | 3<sup>rd</sup> tertile |  |
|  | n | % | n | % | n | % | n | % | n | % | n | % | n | % | n | % | n | % |
| **Type of flooring** |  |  |  |  |  |  |  |  |  |  |  |  |  |  |  |  |  |  |
| Dirt | 50 | 92.6 | 47 | 82.5 | 30 | 54.5 | 56 | 100.0 | 54 | 98.2 | 17 | 30.9 | 55 | 100.0 | 55 | 98.2 | 17 | 30.9 |
| Wood | 0 | 0.0 | 1 | 1.8 | 0 | 0.0 | 0 | 0.0 | 1 | 1.8 | 0 | 0.0 | 0 | 0.0 | 1 | 1.8 | 0 | 0.0 |
| Cement | 4 | 7.4 | 9 | 15.8 | 18 | 32.7 | 0 | 0.0 | 0 | 0.0 | 31 | 56.4 | 0 | 0.0 | 0 | 0.0 | 31 | 56.4 |
| Ceramic | 0 | 0.0 | 0 | 0.0 | 7 | 12.7 | 0 | 0.0 | 0 | 0.0 | 7 | 12.7 | 0 | 0.0 | 0 | 0.0 | 7 | 12.7 |
| **Family bathroom** |  |  |  |  |  |  |  |  |  |  |  |  |  |  |  |  |  |  |
| No | 49 | 90.7 | 50 | 87.7 | 24 | 43.6 | 55 | 98.2 | 48 | 87.3 | 20 | 36.4 | 55 | 100.0 | 49 | 87.5 | 19 | 34.5 |
| Yes | 5 | 9.3 | 7 | 12.3 | 31 | 56.4 | 1 | 1.8 | 7 | 12.7 | 35 | 63.6 | 0 | 0.0 | 7 | 12.5 | 36 | 65.5 |
| **Piped water** |  |  |  |  |  |  |  |  |  |  |  |  |  |  |  |  |  |  |
| No | 31 | 57.4 | 17 | 29.8 | 1 | 1.8 | 32 | 57.1 | 16 | 29.1 | 1 | 1.8 | 33 | 60.0 | 15 | 26.8 | 1 | 1.8 |
| Outside the house | 21 | 38.9 | 36 | 63.2 | 45 | 81.8 | 24 | 42.9 | 38 | 69.1 | 40 | 72.7 | 22 | 40.0 | 40 | 71.4 | 40 | 72.7 |
| Inside the house | 2 | 3.7 | 4 | 7.0 | 9 | 16.4 | 0 | 0.0 | 1 | 1.8 | 14 | 25.5 | 0 | 0.0 | 1 | 1.8 | 14 | 25.5 |
| **Electricity** |  |  |  |  |  |  |  |  |  |  |  |  |  |  |  |  |  |  |
| No | 45 | 83.3 | 27 | 47.4 | 6 | 10.9 | 47 | 83.9 | 27 | 49.1 | 4 | 7.3 | 50 | 90.9 | 24 | 42.9 | 4 | 7.3 |
| Yes | 9 | 16.7 | 30 | 52.6 | 49 | 89.1 | 9 | 16.1 | 28 | 50.9 | 51 | 92.7 | 5 | 9.1 | 32 | 57.1 | 51 | 92.7 |
| **Washing machine** |  |  |  |  |  |  |  |  |  |  |  |  |  |  |  |  |  |  |
| No | 54 | 100.0 | 57 | 100.0 | 27 | 49.1 | 56 | 100.0 | 51 | 92.7 | 31 | 56.4 | 55 | 100.0 | 55 | 98.2 | 28 | 50.9 |
| Yes | 0 | 0.0 | 0 | 0.0 | 28 | 50.9 | 0 | 0.0 | 4 | 7.3 | 24 | 43.6 | 0 | 0.0 | 1 | 1.8 | 27 | 49.1 |
| **Total** | **54** | **100.0** | **57** | **100.0** | **55** | **100.0** | **56** | **100.0** | **55** | **100.0** | **55** | **100.0** | **55** | **100.0** | **56** | **100.0** | **55** | **100.0** |

<sup>1</sup> Based on ownership of durable goods.
<sup>2</sup> Based on characteristics of the household structure.
<sup>3</sup> Based on both ownership of durable goods and characteristics of the household structure.

## DISCUSSION

It is generally acknowledged that the vast majority of the Brazilian indigenous population lives in poverty, although studies assessing and documenting this are scarce.^6^,^11^,^21^ In this setting, and considering the relationship between tuberculosis and poverty, this article presents valuable data from an effort of socioeconomic characterization of what is expected to be an underprivileged subset of an already underprivileged segment of the population: indigenous people with tuberculosis.

Our study didn’t include a control group, as it was not our objective to compare people with and without tuberculosis. However, representative results for the Central-West indigenous population are available in the National Survey and can be used to compare with our findings. The households’ characteristics showed a scenario of lower use of industrial building materials than estimated in the National Survey for the Central-West region, to which Mato Grosso do Sul state belongs. While among our study population 22.9% of households had cement or ceramic floors, 29.5% brick walls and 10.8% clay tile roof, for the National Survey data in the Central-West the proportions of the same categories were 41.7% (95% CI 38.8 to 44.7), 45.3% (95% CI 42.4 to 48.3) and 22.4% (95% CI 19.9 to 24.9),respectively ^11^. According to the authors of the National Survey, the type of building material could be influenced by the access to natural resources on indigenous lands, but would also be related to socioeconomic status.^11^ This was observed in our results. The Polo-Bases with more households made with industrialized materials (Miranda and Aquidauana) were those with less proportion of people with no schooling and higher proportion of households with family bathroom, piped water and electricity. The three calculated SEIs also point in this direction, with most households of Aquidauana and Miranda classified in the 3^rd^ tertile. It is possible that the better socioeconomic conditions found in Aquidauana and Miranda are related to their lower numbers of tuberculosis cases. It is important to remember that all diagnosed cases of tuberculosis among indigenous persons in these Polo-Bases during the study period were included in this study. Despite the small numbers of indigenous cases of tuberculosis in these two Polo-Bases, the results give an indication that those from the south of the state (Amambai and Caarapó) live in poorer socioeconomic conditions than those from the north of the state (Miranda and Aquidauana). In fact, recently it was reported the precarious living conditions in villages in the south of the state, with very rare brick houses and most people living without basic sanitation, very much in line with the results of this study ^22^. The southern part of Mato Grosso do Sul is marked by land conflicts, commonly violent, between the indigenous population and the agricultural sector, the economic base of the state.^23–27^ This violent oppression further aggravates the vulnerability of the indigenous population living there and may worsen their socioeconomic condition. In addition, the border with Paraguay is one of the main routes of entry of drugs in Brazil. Many indigenous people in this region, especially in Amambai, are subjected to intimidation of the organized crime and are recruited to work with the traffic.^28^ The three SEIs showed up as good predictors of socioeconomic status in the study population even though they do not consider any variables that directly reflect income. This was observed when the SEI was analyzed by variables that were not considered in the calculation of the SEIs, but have direct relationship with socioeconomic status (education, regular employment and Bolsa Família). When analyzing by tertiles the distribution of variables included only in two of the three SEIs, similar results were observed. This indicates a potential interchangeability between the three SEIs. Thus, the decision on which SEI to use in a population similar to the one of this study can be based on the availability of the necessary data.

The Combined SEI, that included both durable goods and household characteristics variables, revealed the representativeness of the same variables, with minor changes, as the indexes that analyzed the groups of variables separately. Therefore, even with slightly smaller Kaiser-Meyer-Olkin value than the Household SEI, and less percentage of variance explained (which is expected considering more variables are included), we consider the Combined SEI advantageous as it balances the Goods and the Household SEIs, providing a more robust result. On the other hand, household characteristics may be a result of government actions such as housing programs. In this context, it was expected that the Goods SEI would best represent the purchasing power by including only durable goods, which are not provided by such programs. However, the Goods SEI was the only index that classified people who had regular employment in the 1^st^ tertile, which could indicate less influence of income in this index. Also, the Household SEI showed association with both regular employment and Bolsa Família, indicating that household structure is indeed a good predictor of the socioeconomic status. During fieldwork, in discussions with indigenous people from the communities, it was explained to us that housing programs usually benefit the poorest, but that the poorest people tend to migrate looking for job opportunities. The village leadership then usually passes the empty house to a family of his/her choice.

According to the parameters evaluated, households with more durable goods and made with industrialized materials were considered having a more favorable socioeconomic condition. However, one must consider that this is a classification based on capitalist and consumerist criteria, and may not reflect the actual and/or perceived well-being of the indigenous members. As an example, it is possible that inhabitants of brick houses in a village placed close or inside cities go hungry because they do not have money to buy food, while inhabitants of a house with dirt floor and roof made with thatch can be well nourished with food from the fauna and flora of their land. This is an important limitation and requires caution in interpreting our results. However, we found the indexes here proposed useful in the context of the indigenous peoples living in Mato Grosso do Sul, considering they have had contact with non-indigenous society for more than 200 years and live on small and overcrowded indigenous reservations located in the outskirts of the local town. A different index may be needed for measuring the socioeconomic conditions of indigenous groups in the Amazon region who have maintained their traditional lifestyle, for example.

The proposed socioeconomic characterization incorporates important factors related to health risks, among which stand out the family bathroom, piped water and number of people sleeping in the same room. For tuberculosis transmission, the latter is extremely important considering the potential spread of the disease and that 21.7% of indigenous included in the study reported case of tuberculosis in the family or household in the last two years, and 34.3% more than two years ago (data not shown).

## CONCLUSIONS

The three indexes generated using PCA were useful and were proved suitable for socioeconomic characterization of the study population. Also, it is important to state that the methodology has the potential to be applied in similar indigenous groups, with the advantage of being a simple and consistent approach for socioeconomic characterization in settings were data collection may be difficult. Despite the limitations, this study provides important evidence of the unfavorable living conditions of the indigenous cases of tuberculosis in Mato Grosso do Sul, particularly in areas of land conflict. The low socioeconomic status possibly plays a key role in the high burden of the disease in the region. Future studies comparing the socioeconomic status of indigenous people with and without tuberculosis are of interest.

## ACKNOWLEDGEMENTS

The authors would like to thank the indigenous communities that accepted to be part of this study, the Distrito Sanitário Especial Indígena Mato Grosso do Sul team for their support, and the indigenous healthcare teams for their work and assistance.

